# Fluctuating Environment Can Negate Cheater Success Due to Speed-Agility Trade-Off

**DOI:** 10.1101/2021.06.03.446957

**Authors:** Naomi Iris van den Berg, Lajos Kalmar, Kiran R. Patil

## Abstract

Stability of microbial cooperation through common goods is susceptible to cheating. Evidence suggests that cheating plays a less prominent role in many natural systems than hitherto predicted by models of eco-evolutionary dynamics and evolutionary game theory. While several cheater negating factors such as spatial segregation have been identified, most consider single-nutrient regimes. Here we propose a cheater-suppressing mechanism based on previous experimental observations regarding the biochemical trade-off between growth speed and delay in switching to alternative nutrients. As changing the nutrient source requires redistribution of enzymatic resources to different metabolic pathways, the advantage in speed is offset by lower agility due to longer time required for resource re-allocation. Using an *in silico* model system of sucrose utilisation by *Saccharomyces cerevisiae*, we find that a tradeoff between growth rate and diauxic lag duration can supress cheaters under fluctuating nutrient availability and thereby stabilise cooperation. The resulting temporal dynamics constrain cheaters despite their competitive benefit for the growth on the primary nutrient via avoided public goods synthesis costs. We further show that this speed-agility trade-off can function in synergy with spatial segregation to avoid the collapse of the community due to the cheaters. Taken together, the growth-agility trade-off may contribute to cheater suppression in microbial ecosystems experiencing fluctuating environments, such as plant root microbiota and gut microbiota.

## Introduction

Costs to cooperation provide a selective incentive to defect or cheat (Dumas & Kummerli, 2012). Cheating describes an indirect harm done to non-cheating individuals in a (social) consortium. In contrast to parasites, which incite a direct harm, cheaters act via the intermediary of exploitable social behaviours, the public goods (Ghoul et al., 2014). Cheating cells avoid costs associated with contribution to, but benefit from, the public goods, and hence enjoy a competitive advantage over the cooperative cells. Despite this apparent advantage to the cheaters, cooperation is prevalent in nature, suggesting existence of cheater suppressing mechanisms. Cooperation has been shown to contribute to species and functional diversity; for example, a multi-species community can thrive even on a single carbon source, countering competition in driving ecological dynamics (Blasche et al., 2021; Estrela et al., 2021; Estrela et al., 2020; Goldford et al., 2018). Thus, understanding the mechanisms by which cheating is suppressed and stable cooperation is maintained is of importance in deciphering community structure, dynamics and function.

When unchecked, cheaters in a community can proliferate and disrupt the system’s stability, leading to collapse (Fiegna & Velicer, 2003; Kümmerli et al., 2015; Rainey & Rainey, 2003; Rankin et al., 2007; Ross-Gillespie et al., 2007; Schuster et al., 2017). This extreme outcome would not only require cheater emergence (constrained mostly by stochastic mechanisms), but also cheater proliferation (constrained mostly by non-random, selective mechanisms) (Fiegna & Velicer, 2003). There are three main families of identified mechanisms limiting cheater success. First, inter-group competition, also described as the Simpson’s Paradox, limits the cheater’s relative fitness when assessed from a group selection perspective: within any one group, cheaters outcompete non-cheaters, but groups with more cheaters are outcompeted by groups with relatively fewer cheaters (Chuang et al., 2009; Cremer et al., 2012; van Tatenhove-Pel et al., 2021). Second, cheaters are constrained by relatedness within the community, in accordance with kin(d)-selection theory (Bastiaans et al., 2016; Dennehy & Turner, 2004; Gilbert et al., 2007; Kuzdzal-Fick et al., 2011; Madgwick et al., 2018; Oliveira et al., 2014; Strassmann & Queller, 2011). Third, cheaters can be constrained by mechanisms reducing their access to public goods. Proliferation of cheating mutants may be limited by pleiotropy (Foster et al., 2004), where lower expression of a cooperative trait is itself linked to lower access to a public good. Additionally, grouping together of cooperative cells, facilitated by spatial structuring, can induce a positive, density-dependent, local effect, with cooperative cells passively forming ‘exclusivity zones’ that halt further proliferation of cheating phenotypes (Bachmann et al., 2013; Cavaliere et al., 2017; Folse III & Allison, 2012; Momeni et al., 2013; Nadell et al., 2009; Stump et al., 2018; Van Dyken et al., 2013). This spatial effect can also be made apparent through non-spatially explicit factors such as viscosity of growth media, with higher viscosity favouring cooperative phenotypes by lowering cell dispersal and public goods diffusion (Figueiredo et al., 2021; Kümmerli et al., 2009). Cheater access to public goods can also be actively undermined via enforcement. Enforcement mechanisms reward fellow cooperators and/or punish cheaters (Figueiredo et al., 2021; Ho et al., 2013; Schluter et al., 2015; Smukalla et al., 2008; Travisano & Velicer, 2004). An example of enforcement is that of the rhizobia system in root nodules. If the rhizobia no longer supply fixed nitrogen to their host, the host responds by cutting off supply of carbon exudates to that specific root nodule (Westhoek et al., 2017).

However, these described mechanisms do not always succeed in explaining observed suppressed cheater success. As shown by MacLean and Gudelj (2006), cooperation can persist in homogenous populations, even in the absence of kin(d)-recognition or policing, indicating that other mechanisms are constraining proliferation of cheater cells. Similarly, Dobay et al. (2014) emphasised that interactions between key parameters of public goods cooperation give rise to more complex fitness landscapes, and thus that a further understanding of cooperation in the face of cheating requires multifactorial approaches. Undescribed interactions between multiple cheater-limiting mechanisms (and other (a)biotic processes) may weigh into the many cost-benefit trade-offs involved in microbial community interaction dynamics, and hence alter a cheater’s selective landscape, keeping the role of cheating in natural microbial systems smaller than previously expected (Figueiredo et al., 2021; Waite & Shou, 2012). These additional undescribed and often context-dependent costs thus subject the selective landscape of the cheater strategy to the Cheater’s Dilemma: a set of trade-offs that mitigate the selective benefit imparted by reduced investments in public goods.

In this study, we explore the Cheater’s Dilemma in the context of diauxic shifts. Diauxic shifts are common in microbial ecosystems, yet are underrepresented in microbial community modelling (Pacciani-Mori et al., 2020). Furthermore, in many natural systems, the primary substrate goes through cycles of supply and depletion, such as pulsed carbon additions to plant root microbiota (Berthrong et al., 2013) or pulsed substrate additions (reflecting daily eating patterns) to the gut microbiota (Koch, 1971; Pudlo et al., 2015). Pulsed availability of the primary substrate has been shown to influence community composition and dynamics (Carrero-Colón et al., 2006; Konopka et al., 2007), and may lead to sequential diauxic shifts. These diauxic shifts are characterised by a lag time, primarily due to catabolite repression and transcriptional reprogramming required for re-allocating enzymatic resources between different metabolic pathways (Kremling et al., 2018; New et al., 2014). Trade-offs mediating the diauxic shift lag time have been identified (Chu & Barnes, 2016; Salvy & Hatzimanikatis, 2021; Solopova et al., 2014), where Basan et al. (2020), New et al. (2014) and Fuentes et al. (2021) have showed that faster growing cells experience a longer lag time in shifting to the secondary substrate, whereas slower growing cells resume growth much more quickly. This trade-off falls within an overarching, universal family of trade-offs, based on resource allocation constraints, between (microbial) growth rate and adaptation rate (Kim et al., 2020; Yi & Dean, 2016). Since cheating on public goods production implies faster growth relative to non-cheaters (Ramin & Allison, 2019; Raymond et al., 2012), we here hypothesise that diauxic shifts in a fluctuating environment impose a cost on cheaters. Using a coupled ODE model, we show that the benefit of the cheater strategy depends not only on the relative cost of public good synthesis, but also on the fluctuation regime of the primary substrate and the delay in switching to the secondary substrate. The results bring forward the temporal cost of metabolic switching as an additional mechanism contributing to the Cheater’s Dilemma.

## Model description

To capture the cost to cheating in the context of diauxic shifts, we developed an ordinary differential equations (ODEs) model. We aimed to capture the dynamics of a cooperative microbial species in competition with a subgroup of isogenic cheater mutants. The primary resource in this system is the substrate *S*. We simulate a fluctuating environment with periodic addition of the substrate to a set concentration, *S_max_*. This substrate needs to be hydrolysed into monomers (*M*) in order to become accessible as a carbon/energy source. This is done via extracellular enzymes (*E*) produced by the cooperative ‘producer’ cells (*P*). The monomers form the common goods that the cheaters (*C*) exploit. Both cell types produce a by-product, *B*, during their growth on *M*. When the substrate *S* (and hence *M*) becomes unavailable, both cell-types need to shift to the secondary resource *B*. However, cheaters experience a longer lag time for this shift than the producers, dictated by a constant, *θ*, to account for the speed-agility trade-off (see figure 1 & 2).

**Figure 1.**
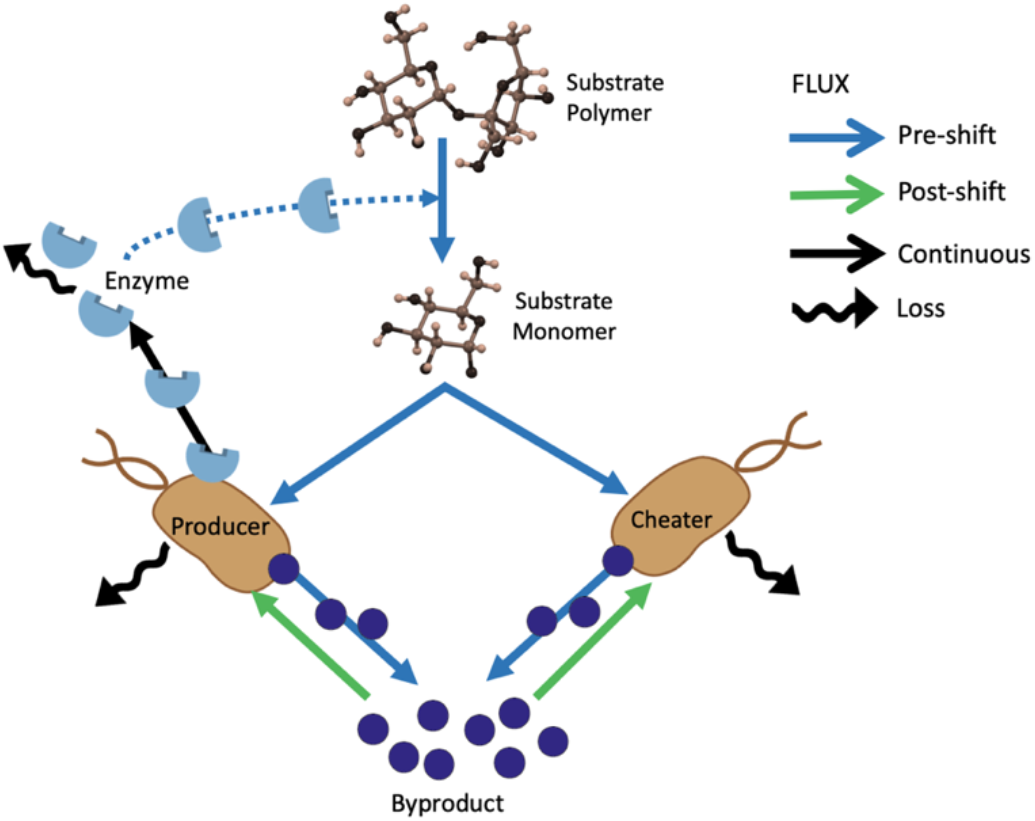
A conceptual diagram of the proposed model. The mass, upon periodic addition as an unavailable polymer, gets bio-transformed as it flows through the system. Some mass gets lost due to thermodynamics and cell mortality (see ‘loss’ arrows). Pre-shift versus postshift fluxes are mutually exclusive (i.e. cannot happen simultaneously), whereas loss and enzyme synthesis fluxes happen continuously (i.e. regardless of growth substrate used).

**Figure 2.**
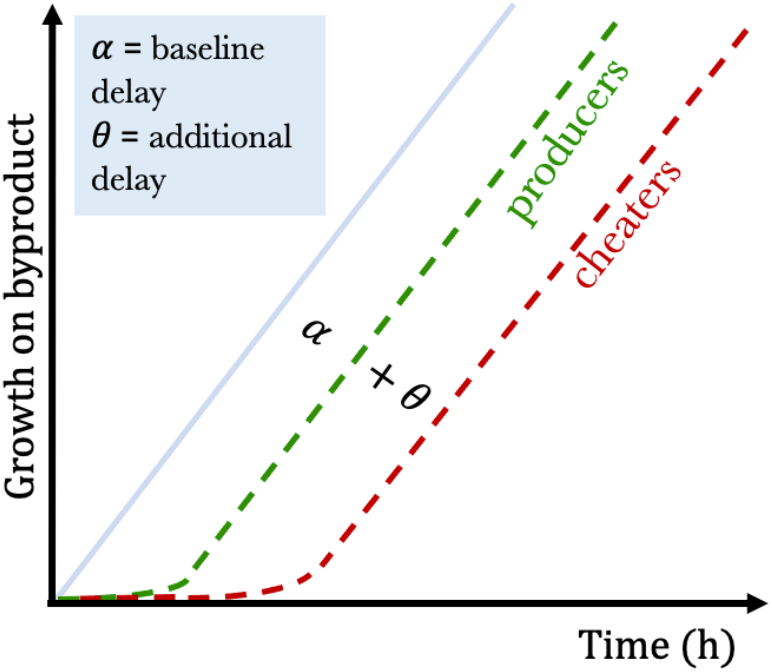
Conceptual graph showing the delay the cheaters and the producers experience when switching to growth on the by-product. The greater the growth rate coefficients, *r_b_* (uptake rate) and *δ_b_* (cell yield), the steeper the growth curves. Without any delay, we assume the per capita uptake rate and therefore growth (since the cell yield coefficient is constant) is a linear function of by-product availability. With delay, we assume an expo-linear growth curve instead, where the maximum linear per capita growth rate is initiated after a determined delay period (Goudriaan & Monteith, 1990). The bigger *α*, the longer the delay for growth on byproduct. For cheaters, this delay is longer (dictated by magnitude of *θ*).

A concrete example of a microbial system resembling the model explored here is sucrose utilisation by the yeast *Saccharomyces cerevisiae*. This model microbe has a long history of use in food and drink fermentations, and is also widely used in biotechnological applications such as production of biofuels and pharmaceuticals (Marques et al., 2016). *Saccharomyces cerevisiae* secretes invertase enzyme (β-D-fructofuranosidase, E.C. 3.2.1.26) to hydrolyse sucrose. Ethanol is excreted as a by-product during growth, which can later be consumed when the hydrolysed monosaccharides (i.e. glucose and/or fructose) are exhausted (Atiyeh & Duvnjak, 2002; Gibb, 2018). The change from growth on sugars to growth on ethanol constitutes the diauxic shift.

While the hydrolysis reaction via invertase is performed in the periplasmic space, it has been demonstrated that most (~99%) of the monosaccharides released are extracellular and need to be imported (Basso et al., 2011; Gore et al., 2009). Hence, invertase can, for the purpose of this study, be considered functionally equivalent to purely extracellular enzymes, as done by the previous studies looking into cheater dynamics in *S. cerevisiae* (Cavaliere et al., 2017; Chen et al., 2014; Gore et al., 2009; Greig & Travisano, 2004; Koschwanez et al., 2011; Travisano & Velicer, 2004; Van Dyken et al., 2013). The impact of pulsed carbon additions on *S. cerevisiae* diauxic shift and growth has been studied extensively (Espinosa Vidal et al., 2015; Ronen & Botstein, 2006; van den Brink et al., 2008), however, not in the context of cheater dynamics and the speed-agility trade-off.

In our model, we assume that the upper limit of glucose concentration leading to repression of invertase synthesis (as identified by Dueñas-Sánchez et al. (2012)) will not be reached due to the slow release and immediate uptake of glucose in our system. With ethanol as our byproduct, and periodic minimal additions of primary substrate, we assume its concentrations do not reach toxicity levels of >10%. Local structure allows the grouping together of cells, which decreases the likelihood of monomers diffusing too far away for consumption by producers, benefitting producer growth. This locality benefit does not apply for by-product uptake, since the by-products may diffuse away further as they are not used immediately (i.e. >1 doubling times between production and consumption of by-product). We assume this local effect proportionally increases producer uptake (and therefore growth) rates. For universal model assumptions, see table 1.

**Table 1.** Overview of model assumptions

| Assumption | Model implications | Justification |
| --- | --- | --- |
| We simplify uptake rates by assuming linear dynamics as a function of monomer/by-product concentration instead of Monod dynamics. In this linearised realm of uptake, only by-product uptake rate operates near maximum velocity. | $r_b \approx r_{b_{max}}$<br>$r_b \ll r_M < r_{M_{max}}$ | |
| Both cheaters and producers experience the same linear per capita uptake rate of the by-product, which commences after an initial ‘baseline’ delay. However, cheaters experience an additional delay since they grow faster on the primary substrate. | Cheater total delay duration is described by $\alpha + \theta$ (see figure 2) | Basan et al. (2020); New et al. (2014), Fuentes et al. (2021), Raymond et al. (2012), Ramin and Allison (2019) |
| Growth on the primary substrate continues until its concentration reaches virtually 0, after which the diauxic shift commences | $r_M = 0$ when $M < M_{threshold}$ | Haurie et al. (2003) |
| By-products continue to accumulate in the system until they start to be consumed. Consumption halts upon exhaustion. | $r_b = 0$ when $M > M_{threshold}$ or $B < B_{threshold}$ | Haurie et al. (2003) |
| We assume no conversion of substrate into processed monomer when the enzyme or the substrate is missing. | Input $M$ only via Michaelis-Menten decay of $S$. | Schuster et al. (2017) |
| Cheater mutants cannot regain enzyme-synthesising abilities. | $\gamma=0$ for $C$ throughout simulated time | |
| The system is constrained by an explicit carrying capacity, implying competition for space or other limiting resources. | Terms related to mass uptake and secretion are constrained as $(C + P) \rightarrow K$ | Frank (2010); Smith et al. (2014) |
| Local structure allows the grouping together of cells, which decreases the likelihood of monomers diffusing too far away for consumption by producers, benefitting producer growth. | A proportionality constant of $\varphi > 1$ increases $M$ uptake term for producers. | Stump et al. (2018) |
| The environment (e.g. temperature and pH), other than the dynamic availability of the primary substrate, is held constant. | Kinetic parameters, $V_{max}$ & $K_m$, are constant | |
| Cheater mutants are established and have survived stochastic extinction. They initially occur in small numbers in an otherwise stable population of producer cells, to reflect a realistic order of magnitude of early mutant cell line evolution. | The model is initialised with a cheater to producer ratio of $\leq 1:10$ | Ross-Gillespie et al. (2007) |

The following coupled ODEs describe the dynamics in our system:

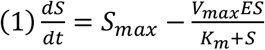

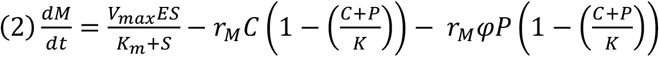

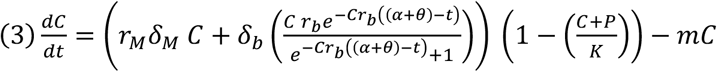

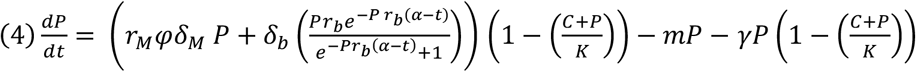

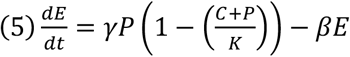

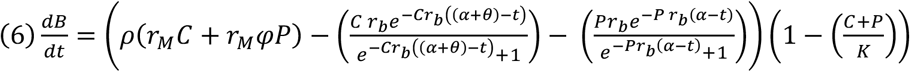

## Model analysis

Assuming constant primary substrate and monomer availability (i.e. no diauxic shift), and no carrying capacity, our system’s Jacobian can be reduced to:

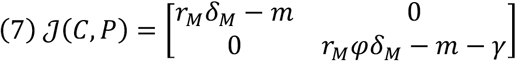

This leads to the following condition, exclusively reliant on a local benefit effect, in which producers cannot be outcompeted by cheaters (with all parameters >0 and *τ_M_, δ_M_* & *m* being equal for both cell types):

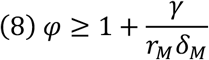

To investigate the effect of substrate availability dynamics, we performed numerical simulation experiments on the model with constant parameter values (see Methods). We performed two modelling experiments to identify qualitative behaviour of cheater-producer dynamics under serial batch culture conditions, replacing half of the volume with fresh medium at each addition event (simulating periodicities where dilution is not faster than growth). In the first model experiment, we simulate along a range of periodicity and additional delay values, and we assume there to be no local effect that benefits producers (*φ* = 1). In the second simulation experiment, we additionally consider that the spatial/structural heterogeneity in the environment is such that producers experience local benefit effect (*φ* > 1). A local sensitivity analysis was executed for parameters within their defined biological range (see table 2) for various periodicity regimes (6h, 12h, 24h), assuming intermediate strength of the local benefit effect (*φ* = 1.3) and estimated additional delay (*θ* = 3). The sensitivity was measured for time until the take-over event (where *C* > *P*). Since the model output can represent three qualitatively different trajectories, namely: 1) take-over followed by system collapse, 2) no take-over and continued producer stability (i.e. stable cheater suppression, and 3) no takeover and system collapse (i.e. extinction of producer before cheater can take over), we defined the local sensitivity of our model to each of the parameters by assessing whether the model output is qualitatively different between the two extremities of the defined parameter value range. The sensitivity analysis results shows that the sensitivity of the model output to each individual parameter is primarily dependent on the primary substrate regime. For example, in long periodicities (24h), producers benefit from a higher mortality rate, whereas in intermediate periodicities (12h) this same mortality rate leads to cheater take-over and collapse (Table 3).

**Table 2.** Overview of parameters used in our model

| Parameter | Description | Dimensions | Value | Justification |
| --- | --- | --- | --- | --- |
| $S_{max}$ | Max amount of substrate added at each pulse | $[g\ l^{-1}][h]^{-1}$ | 20 | Corresponding to standard medium enrichment of 2% sucrose. Francesca et al. (2010); Van Dyken et al. (2013). |
| $V_{max}$ | Maximum reaction rate enzyme with substrate | $[g\ l^{-1}][h]^{-1}$ | ~20.5 | For soluble invertase hydrolysing sucrose at optimum pH and temperature, adopted from Essel and Osei (2014). Original units: 1 mM min<sup>-1</sup>. |
| $K_m$ | Half-saturation constant for substrate decay | $[g\ l^{-1}]$ | ~8.2 | For soluble invertase hydrolysing sucrose at optimum pH and temperature, adopted from Essel and Osei (2014). Original units: 24 mM. |
| $r_M$ | Uptake rate primary monomer | $[g\ l^{-1}][h]^{-1}$ | ~2 | Assuming a less than maximum growth rate, e.g. 0.3 h<sup>-1</sup>, the glucose uptake rate becomes ~2 h<sup>-1</sup>. Jones and Kompala (1999). |
| $r_b$ | Uptake rate by-product | $[g\ l^{-1}][h]^{-1}$ | ~0.3 | Ethanol uptake rate is around ~0.3 h<sup>-1</sup> (with max growth rate of 0.2 h<sup>-1</sup> and cell yield of 0.65). Jones and Kompala (1999). |
| $\delta_M$ | Yield coefficient primary monomer | $[g\ l^{-1}]^{-1}$ | ~0.15 | *S. cerevisiae* growing on glucose. Jones and Kompala (1999). |
| $\delta_b$ | Yield coefficient by-product | $[g\ l^{-1}]^{-1}$ | ~0.65 | *S. cerevisiae* growing on ethanol. Jones and Kompala (1999). |
| $\varphi$ | Proportionality constant for local benefit in producer growth | Dimension-less | 1.3±0.2 | Based on empirical evidence of positive assortment of cooperative yeast cells, adopted from Momeni et al. (2013). |
| $\rho$ | By-product production/excretion | $[g\ l^{-1}]^{-1}$ | ~0.4 – 0.5 | Around half (40-50%) of each gram of glucose taken up is turned into ethanol, whereas 15% goes to biomass production (see $\delta_M$). The rest is lost as heat, CO<sub>2</sub>, etc. Taherzadeh et al. (1997). |
| $m$ | Self-maintenance costs & cell death | $[h]^{-1}$ | 0.05 ± 0.02 | Adopted from Momeni et al. (2013) and Shou et al. (2007). |
| $\gamma$ | Enzyme production | $[h]^{-1}$ | 0.03 ± 0.02 | 1 – 5 % fitness advantage by cheaters. Giuseppin et al. (1993), Christiansen and Nielsen (2002), Gore et al. (2009), Momeni et al. (2013), Waite and Shou (2012). |
| $\alpha$ | Amount of delay in expo-linear growth on by-product | $[h]$ | ~3 | Yeast cells experience a lag time of 3 to 4 hours to shift from glucose to ethanol. Lewis et al. (1993), Breining and Jespersen (2002), Haurie et al. (2003). |
| $\beta$ | Enzyme loss | $[h]^{-1}$ | 0.3±0.2 | 10 – 50% of enzymes decay or become inactive every hour. Bassetti et al. (2000); Folse III and Allison (2012). |
| $\theta$ | Amount of additional delay in expo-linear growth on by-product | $[h]$ | 3±1 | In accordance with the additional delay of 3±1 hours in the shift from glycerol to acetate by mutant bacterium with higher glycerol uptake as demonstrated by Basan et al. (2020). Same study assumes similar trade-off magnitude for yeast. |
| $K$ | Carrying capacity | $[g\ l^{-1}]$ | ~30 | Empirically shown to range from 20 to 55. Malairuang et al. (2020). |

**Table 3.** Local sensitivity analysis assuming a local benefit effect and additional delay in serial batch culture conditions. Qualitative trajectories model output for minimum and maximum value of parameter range: 1) take-over followed by system collapse (blue shade), 2) no take-over and continued producer stability (i.e. stable cheater suppression, yellow shade), and 3) no take-over and system collapse (i.e. extinction of producer before cheater can take over, grey shade).

| Parameter | Minimum value | Maximum value | Default | Periodicity | Output for minimum value parameter | Output for maximum value parameter |
| --- | --- | --- | --- | --- | --- | --- |
| $\rho$ | 0.4 | 0.5 | 0.5 | 6 | 2 | 2 |
| $m$ | 0.03 | 0.07 | 0.03 | 6 | 2 | 2 |
| $\gamma$ | 0.01 | 0.05 | 0.05 | 6 | 3 | 2 |
| $\alpha$ | 3 | 4 | 3 | 6 | 2 | 2 |
| $\beta$ | 0.1 | 0.5 | 0.2 | 6 | 2 | 3 |
| $K$ | 20 | 55 | 30 | 6 | 2 | 2 |
| $\rho$ | 0.4 | 0.5 | 0.5 | 12 | 2 | 2 |
| $m$ | 0.03 | 0.07 | 0.03 | 12 | 2 | 1 |
| $\gamma$ | 0.01 | 0.05 | 0.05 | 12 | 3 | 2 |
| $\alpha$ | 3 | 4 | 3 | 12 | 2 | 2 |
| $\beta$ | 0.1 | 0.5 | 0.2 | 12 | 2 | 1 |
| $K$ | 20 | 55 | 30 | 12 | 2 | 2 |
| $\rho$ | 0.4 | 0.5 | 0.5 | 24 | 1 | 1 |
| $m$ | 0.03 | 0.07 | 0.03 | 24 | 1 | 2 |
| $\gamma$ | 0.01 | 0.05 | 0.05 | 24 | 2 | 1 |
| $\alpha$ | 3 | 4 | 3 | 24 | 1 | 1 |
| $\beta$ | 0.1 | 0.5 | 0.2 | 24 | 1 | 1 |
| $K$ | 20 | 55 | 30 | 24 | 1 | 1 |

## Results

In the absence of a speed-agility trade-off faced by cheater cells (*θ* = 0), and/or a local benefit enjoyed by cooperative producer cells (*φ* = 1), a small subgroup of cheater mutant cells is bound to outcompete producer cells, regardless of periodicity of primary substrate (see panel A in figure 4). Collapse of the system follows after cheaters take over, which is in line with *in silico* (Szilágyi et al., 2017), as well as *in vitro* findings (Allison et al., 2014). Figure 3 shows a simulation example of such a takeover event.

**Figure 3.**
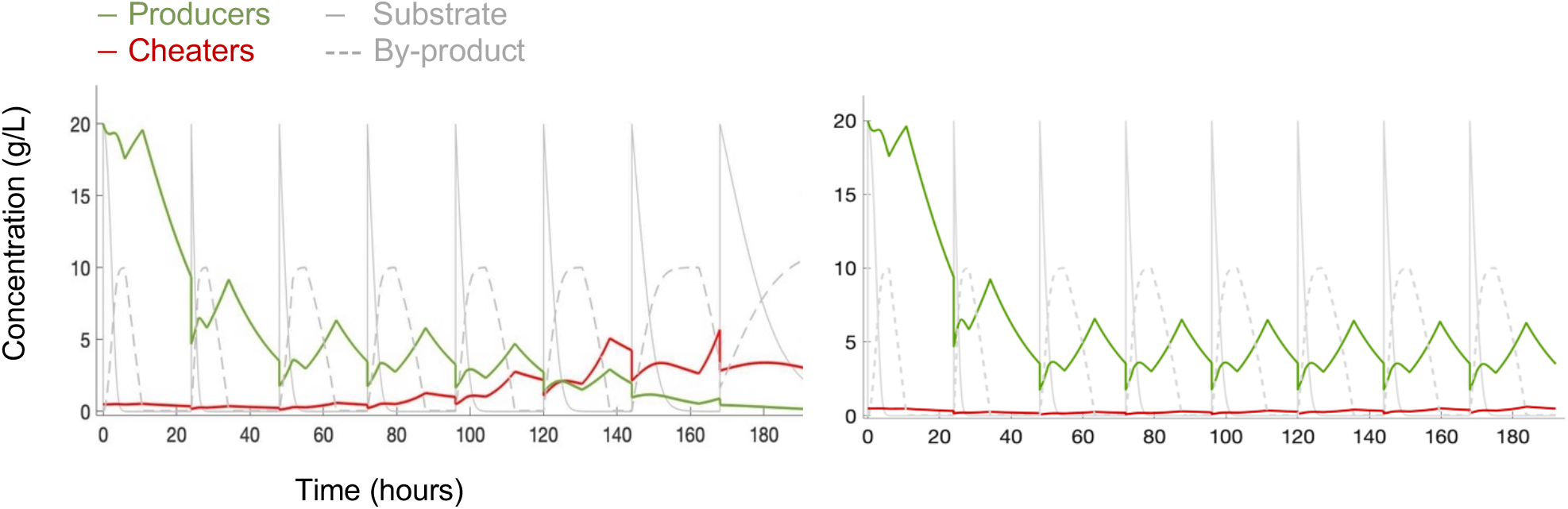
Left: simulation without any cheater-negating mechanisms. Cheaters take over after a few substrate additions (in intervals of 24h), followed by system collapse. Right: example of system stability with cheater-negating mechanisms switched on. Cheaters do not take over, and producer populations remain stable.

**Figure 4.**
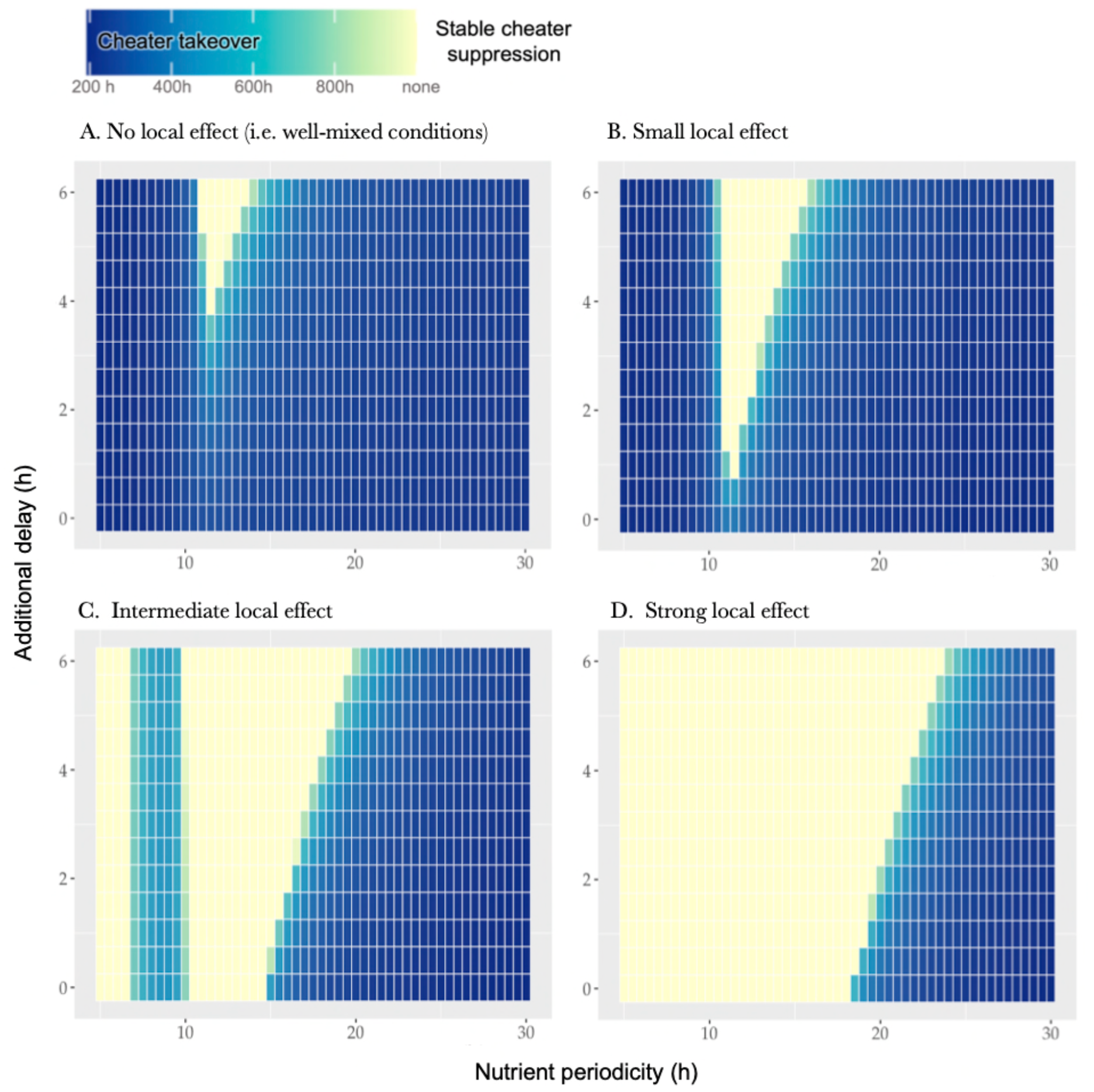
Model solutions along a range of periodicity (5 – 30h) and additional delay values (0 – 6h). Solutions predict a realm of system stability (i.e. cheater suppression) along periodicity regimes, which expands with the strength of the speed-agility trade-off (i.e. the length of additional delay) and strength of the local benefit effect (which increases from panel A to D).

Yet, with an additional delay experienced by the cheaters and/or with a local benefit effect for producers, cheater proliferation is no longer the inevitable outcome. The model output predicts the presence of a realm of system stability, which expands with length of additional delay and strength of the local benefit effect, where the pulse interval is neither too short nor too long. When too short, a diauxic shift never occurs. In this realm, cheaters can exercise their competitive benefit over the primary substrate continuously, without suffering the consequences of a delayed growth on the secondary substrate (rendering the amount of additional delay irrelevant). However, the local benefit effect, if available, can erode the cheater’s competitive overhand in this regime.

When the period between substrate additions is too long, both substrates can get exhausted within one cycle, after which the producers can no longer offset the costs of producing enzymes with the benefit in growth experienced on the by-product, leading to a realm where cheaters outcompete producers again. Yet, in the intermediate periodicity regimes, an additional delay in combination with a local benefit effect can maintain system stability and supress cheaters from taking over. In this regime, either mechanism alone would lead to collapse suggesting a synergy between the two cheater-negating mechanisms.

## Discussion & Conclusion

In this study, we aimed to elucidate *in silico* that the selective benefit of the cheater strategy in microbial public goods systems depends not only on the cheater’s relative advantage over cooperators (i.e. the costs associated with the production of the public good that cheaters defect on), but also on (spatio-) temporal characteristics of the system, subjecting cells that are defecting or about to defect to a more complex selective landscape limiting their real-life occurrence: the ‘Cheater’s Dilemma’. More specifically, we investigated the Cheater’s Dilemma in the context of diauxic shifts. We examined the impact of the availability dynamics of the primary substrate (that is hydrolysed and made available by the public good) on the cheater’s ability to outcompete non-cheaters, taking into account two cheater-proliferation negating mechanisms: a speed-agility trade-off for cheater cells – which is grounded on first principles and recent experimental observations (Basan et al., 2020; Fuentes et al., 2021; New et al., 2014) – and a local growth benefit for cooperative (producer) cells.

Our mathematical model suggests that a speed-agility trade-off can make an appreciable difference to the community stability in the wake of cheaters. The identified temporal complexity, namely the interaction between the nutrient periodicity regime and the speed-agility trade-off, may constrain cheater mutants from proliferating regardless of their competitive benefit, which would be overlooked in a single regime of substrate availability and diauxic delay.

The proposed mechanism of the speed-agility trade-off can work synergistically with other cheater-negating mechanisms; for example with the local benefit effect, as we showed. This synergy can expand the range of conditions under which the system is stable wherein either mechanism alone would lead to system collapse. Accounting for multiple cheater-negating mechanisms will therefore be important to identify critical transitions in fluctuating environments with cheater emergence (Scheffer et al., 2012). Since our model is not spatially explicit, we incorporated the local benefit effect via the simplified alternative of a parameter. However, as argued by Allen et al. (2013), such a parameter is limited in the extent to which it is reflective of true (and variable) local effects experienced by cells, such as colony geometry and spatial diffusion. For the purpose of this study – namely qualitatively demonstrating whether and to which extent the two studied mechanisms have the potential to limit cheater proliferation in the context of diauxic shifts and substrate periodicity – we consider the simplification to be justified.

A possible limitation to our model is that it assumes periodicity of substrate additions that is fixed and not at all stochastic, which could have evolutionary consequences (see Levins (2020); Mitchell et al. (2009)), for instance in duration of lag phase (Chu & Barnes, 2016). However, for many natural systems, such pulsation of carbon additions is regular and at most stochastic within a limited range of this regularity, and hence, one may expect the possible role of stochasticity to be of limited relevance to the predicted qualitative behaviour of the modelled cheater – producer dynamics. Having modelled such stochasticity (by drawing a random number within a defined range for each substrate addition), the qualitative behaviour of the model remains practically identical (see supplement C - E).

In the realm of longer pulse intervals, where the speed-agility trade-off comes into effect, producer stability is maintained via a process akin to the Snowdrift game, facilitating coexistence rather than full extinction of the cheater cells (in line with Xenophontos et al. (2021)). In this realm, shorter diauxic delay times provide a selective benefit to producer cells. Additionally, in the realm of very long pulse intervals (with imminent substrate exhaustion), one may expect a selective pressure towards evolving optimality in regulation of enzyme production, not only in the (empirically identified) upper limit of substrate concentration (Dueñas-Sánchez et al., 2012), but also in the lower limit, allowing producers a level playing field in periods of starvation. This lower limit of the primary substrate concentration could also serve as a signal in asymmetric anticipatory regulation of the diauxic shift, optimising the delay period in periodicity realms where pulsations are long (Mitchell et al., 2009). Having modelled such conditional enzyme investment, we still see similar qualitative behaviour of our model, generating a realm of stability that expands with the additional delay period (see supplement A).

In conclusion, our results highlight the need to study cheater dynamics in a more complex (temporal) dynamic landscape, and call for future studies (*in vitro* and *in silico* alike) to take nutrient periodicity and the speed-agility trade-off into account when mapping the potential proliferation of (microbial) cheater mutants in fluctuating environments, such as the gut.

## Methods

Numerical simulations were executed using the ‘ode45’ function in MATLAB (R2021a). Heatmaps of model output were generated using the ‘ggplot2’ package in RStudio (Version 1.3.1093).

## Acknowledgements

We thank Anna Lindell for providing parts of the illustrations used in this study. This project has received funding from the European Research Council (ERC) under the European Union’s Horizon 2020 research and innovation programme (Grant agreement No. 866028).

## Author contribution

N.I.v.d.B. and K.R.P. conceived the project and wrote the manuscript. N.I.v.d.B. and L.K. performed the model simulations. All contributed to data interpretation.

---

**Supplementary Figures.**
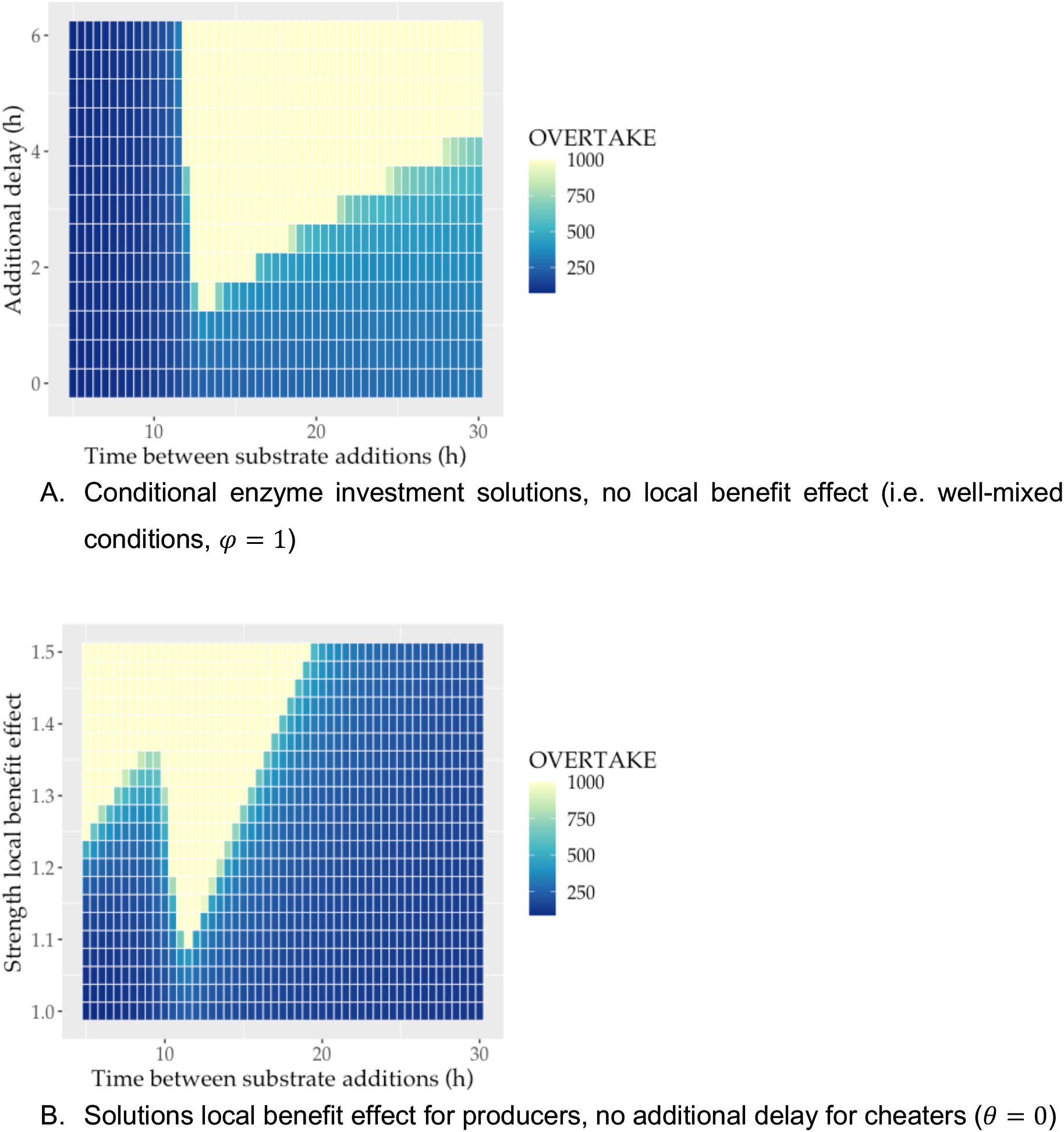
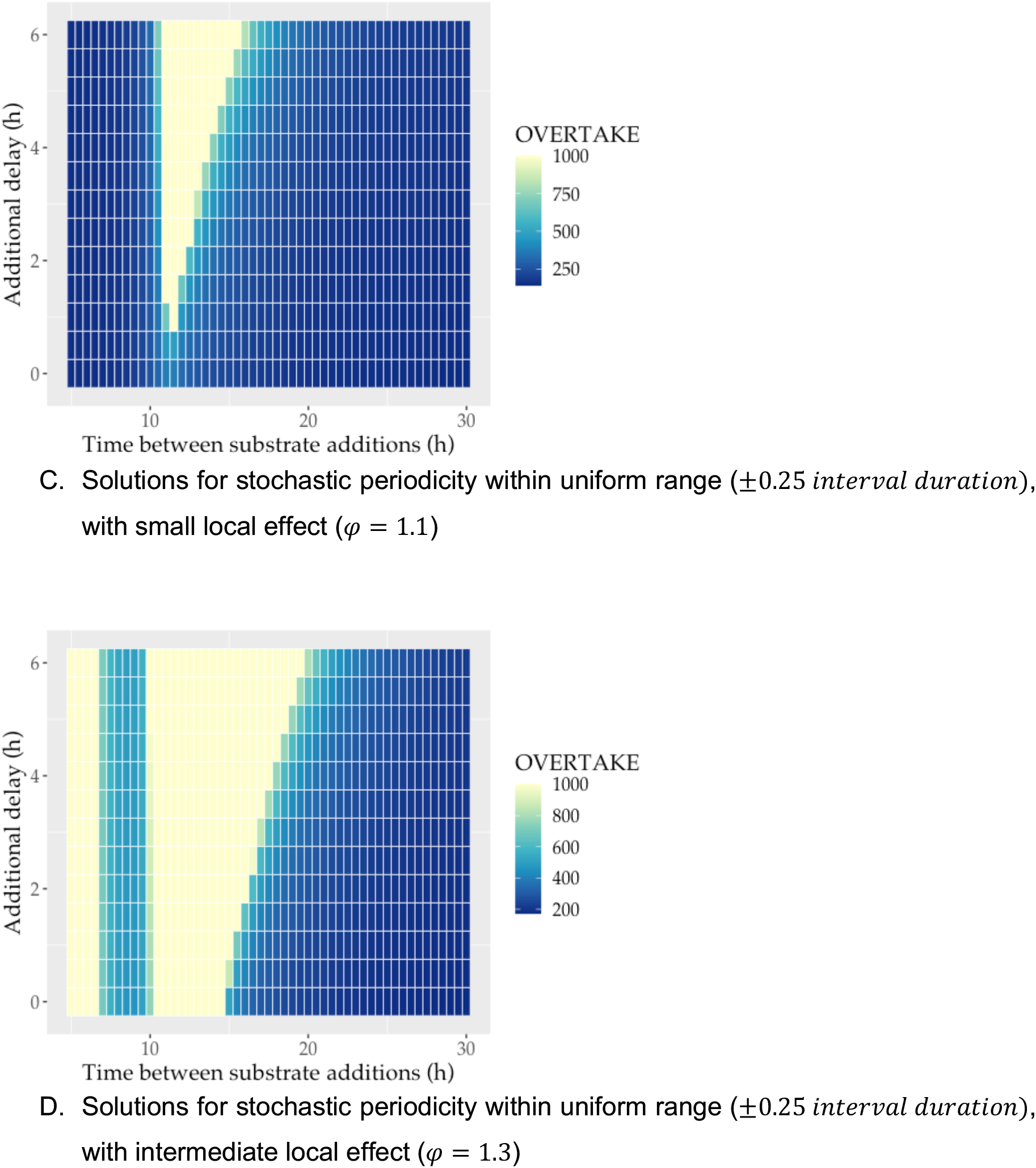
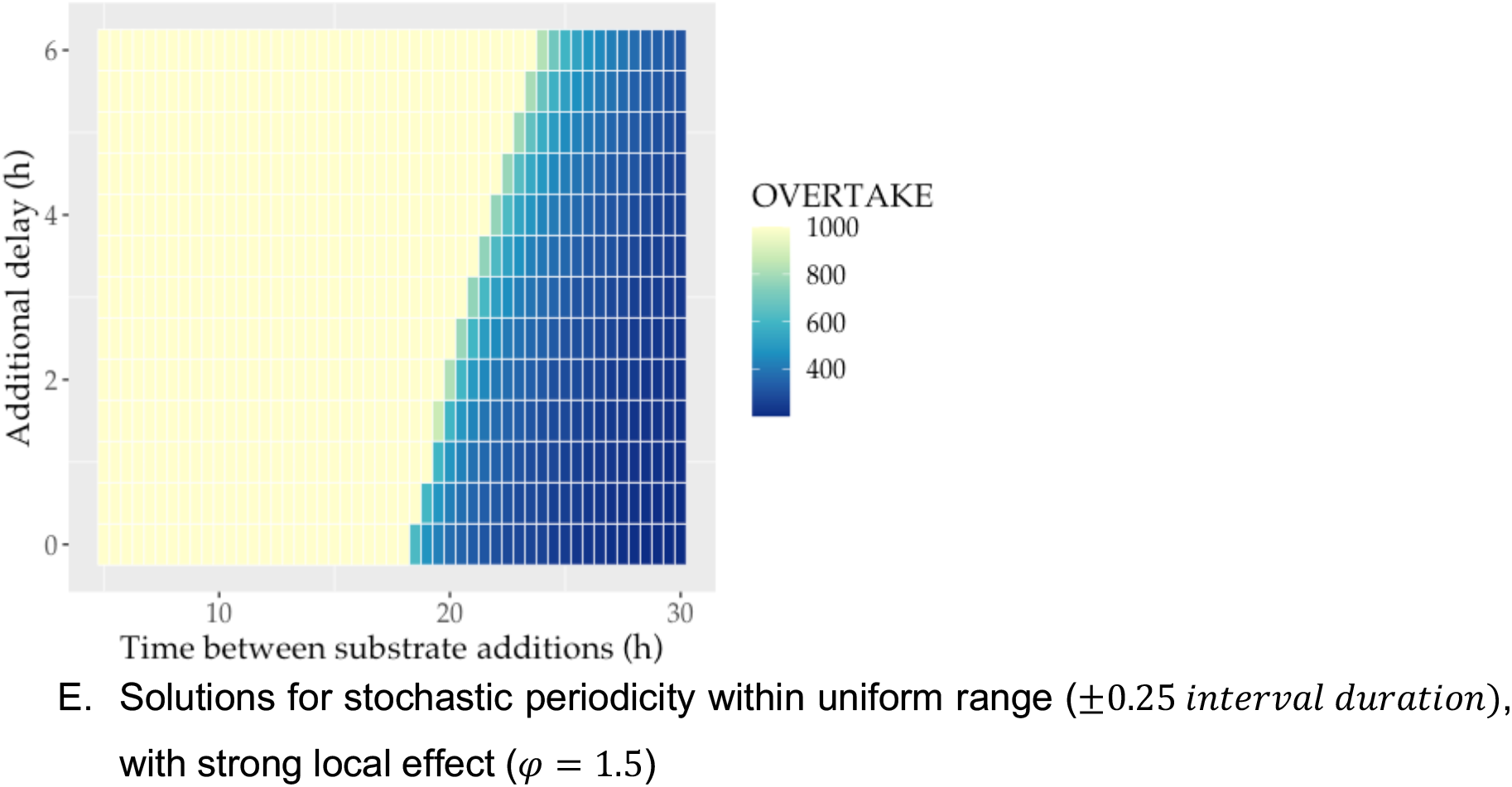

## Notes

### Competing Interest Statement

The authors have declared no competing interest.

